# Genetic stock identification reveals greater use of an oceanic feeding ground around the Faroe Islands by multi-sea winter Atlantic salmon

**DOI:** 10.1101/2022.07.04.498682

**Authors:** Ronan James O’Sullivan, Mikhail Ozerov, Geir H. Bolstad, John Gilbey, Jan Arge Jacobsen, Jaakko Erkinaro, Audun H. Rikardsen, Kjetil Hindar, Tutku Aykanat

**Affiliations:** Organismal and Evolutionary Biology Research Program, Faculty of Biological and Environmental Sciences, PO Box 56, FI-00014 University of Helsinki, Finland; Biodiversity Unit, University of Turku, FI-20014 Turku, Finland; Norwegian Institute for Nature Research (NINA), NO-7485 Trondheim, Norway; Marine Scotland Science, Freshwater Fisheries Laboratory, Faskally, Pitlochry, Scotland PH16 5LB, UK; Faroe Marine Research Institute, Nóatún 1, FO-100 Tórshavn, Faroe Islands; Natural Resources Institute Finland (Luke), POB 413, FI-90014 Oulu, Finland; Department of Arctic and Marine Biology, UiT The Arctic University of Norway, NO-9037 Tromsø, Norway

**Keywords:** *Salmo salar*, spatial variation in resource use, age class structure, phenotypic diversity, migration, Faroe Islands

## Abstract

There is a general paucity of knowledge regards spatial variation in marine resource use for many taxa, even those of high socio-economic importance such as Atlantic salmon. While it is known that the oceans around the Faroe Islands support a salmon feeding ground, the relative use of this feeding ground by different age classes across different stocks remains largely unexplored. Using a combination of genetic stock assignment and run-reconstruction models, we observed a consistent pattern whereby the proportion of multi-sea winter (MSW) salmon for a given reporting group was substantially greater around the Faroes than the MSW proportion for that reporting groups among the pre-fisheries abundance. Surprisingly, MSW fish from Ireland and UK were as likely to occur around the Faroes as were MSW fish from more north-eastern regions such as the Teno river and the Barents and White Seas. MSW fish from Southern Norway were the most likely to be caught at the Faroes. While 1SW salmon from Ireland and UK as well as from Southern Norway occurred at similar rates around the Faroes, 1SW fish from more north-eastern reporting groups were nearly entirely absent from the same feeding ground. In combination with previous studies that examine the marine distribution of Atlantic salmon, our results strongly indicate that the oceans around the Faroes play host to a predominantly MSW salmon feeding ground and that use of this resource varies both within the age classes of a given stock as well as between different stocks. Furthermore, these results suggest that MSW fish from certain stocks might preferentially undertake migrations to the Faroes. Variation in spatial resource use may help to buffer salmon stocks against localised negative changes in marine conditions.

## Introduction

Understanding patterns of resource use provides insight into the evolution and maintenance of the intraspecific phenotypic diversity observed within many aquatic taxa (Hawley et al. 2016; Kang and Thibert-Plante 2017; Kane et al. 2022). Differential resource use can limit intraspecific competition across life stages as well as buffer against extinction risk by spatially segregating a population into different habitats (Schindler et al. 2015; Østbye et al. 2020). From an applied perspective, knowledge of which and when individuals in a population exploit a given resource can be used to determine the efficacy of marine protected areas (Harada et al. 2015; Hernández et al. 2019), regulate fisheries (Armstrong et al. 2013), and generate anticipatory predictions to help inform policy (Eikeset et al. 2013; Ayllón et al. 2018, 2019). However, the ecology of resource use remains poorly understood for many taxa with oceanic life stages. One such taxon where many aspects of the marine ecology are poorly understood is Atlantic salmon, *Salmo salar*.

The marine ecology of Atlantic salmon was, until relatively recently, almost completely unknown. There has been a sustained effort at determining migration routes (Hansen & Jacobsen, 2003; Spares et al. 2007; Strøm et al. 2017, 2018; Bradbury et al. 2021; Gilbey et al. 2021; Rikardsen et al. 2021), dietary composition (Jacobsen & Hansen 2001; Rikardsen & Dempson 2011; Utne et al. 2022), disease transmission (Teffer et al. 2020), oceanic predation (Strøm et al. 2019), and the stocks from which salmon caught at sea originate (Bradbury et al. 2016, 2021; Gilbey et al. 2017; Ó Maoiléidigh et al. 2018). This increase in research effort has coincided with a realisation that changes in the oceanic environment have precipitated declines in the marine survival rate and abundance of Atlantic salmon (ICES 2021; Thorstad et al. 2021). For example, a reduction in their growth at sea (Vollset et al. 2022) and a deterioration in marine feeding conditions (Utne et al. 2021) have both been observed. Czorlich et al. (2022) demonstrated that anthropogenic exploitation of capelin, *Mallotus villosus*, stocks in the Barents Sea has induced evolution towards earlier maturation in salmon from the Teno. Together, these studies suggest that the decrease in survival/abundance of Atlantic salmon is influenced by the availability of oceanic food resources and that human-induced changes are unequivocally responsible.

Though purely freshwater resident populations are known (Hutchings et al. 2019), the majority of Atlantic salmon populations are anadromous. Anadromous salmon hatch in freshwater where they spend one to several years before undertaking an oceanic feeding migration. In most cases, this migration lasts between one and four years (Thorstad et al. 2011) after which the fish returns to breed in its natal river (although straying and breeding in non-natal rivers can occur [Keefer & Caudill, 2014]). Salmon that spend a single year at sea are referred to as ‘one-sea winter’ fish, and those spending longer known as ‘multi-sea winter’. Marine feeding opportunities are far greater and more nutritious than those found in freshwater (Gross et al. 1988). As such, when anadromous salmon return to spawn, they are larger and, thus, more fecund than they would be if they had remained in freshwater (though the costs and benefits of anadromy differ between sexes, see Fleming 1998). These potential fitness benefits are balanced against far higher mortality rates in the marine environment, i.e. due to higher predation/starvation/exploitation rates at sea (Aas et al. 2011). For evolved behaviours such as marine feeding migrations to remain adaptive, the anadromous portions of Atlantic salmon stocks must experience relatively invariant oceanic conditions across multiple generations. Thus, disruption to the spatio-temporal availability of marine prey such that migrations no longer coincide with peaks in resource availability could negatively impact Atlantic salmon stocks in much the way as has already been observed (ICES 2021; Utne et al. 2021; Vollset et al. 2022). If resource use varies spatially across age classes within a stock, this likely exposes that stock to a larger suite of stressors than if resource use patterns were uniform across age (cf. Schindler et al. 2015). As a corollary, variation in resource use between stocks could make some more susceptible to the deleterious effects of anthropogenic change than others.

How oceanic resource use varies spatially across age classes of Atlantic salmon both within and between stocks remains largely unexplored (Rikardsen & Dempson, 2011). Quantifying this spatial variation has the potential to inform discussion as to which age classes and stocks are most likely to be impacted by spatio-temporally dependent changes in marine conditions and resource availability. To investigate this topic, Atlantic salmon were sampled at one of their known feeding grounds around the Faroe Islands (Fig. 1; Fig. S1). Sampled fish were assigned to their reporting group of origin using a geographically broad genetic baseline of 50 populations ranging from the Pechora in the Russian Federation to the east coast of North America. Coupled with scale-read sea ages, we determined the proportion of multi-sea winter salmon from each reporting group represented in the fishery. Run-reconstruction was then used to provide a quantitative comparison of the proportions of multi-sea winter salmon caught around the Faroes to the proportion of multi-sea winter salmon for each reporting group among deep-sea pre-fisheries abundance.

**Figure 1:**
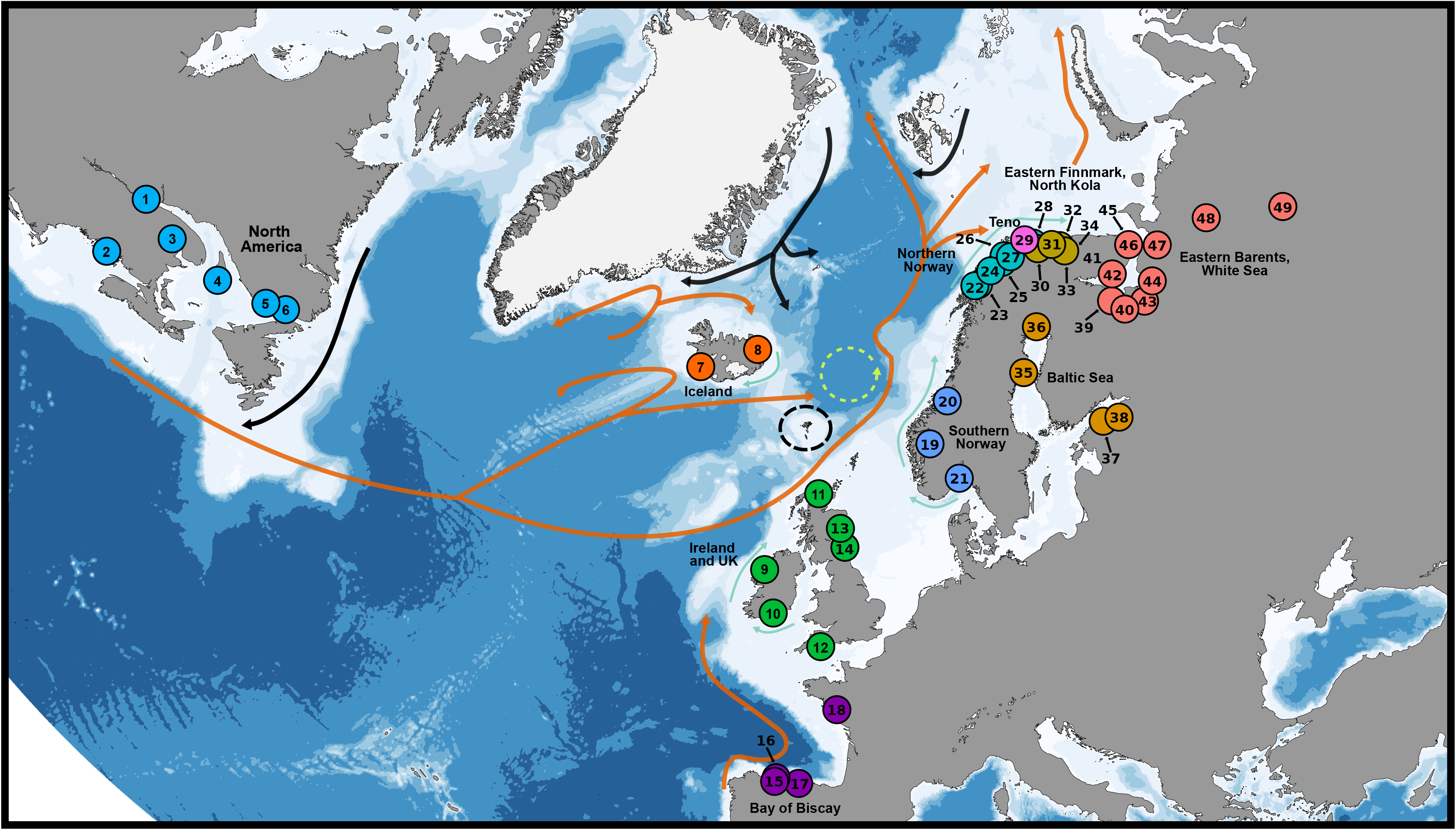
Agglomerative reporting groups of Atlantic salmon (bold, black text) and the populations in each reporting group used in the genetic baseline. Numbers correspond to population names in Tables S1 and S3. Warm Atlantic currents are denoted using orange arrows, cold polar currents from the Arctic Ocean using black arrows, and coastal currents using turquoise arrows. The Vøring Gyre and its direction of flow are denoted using the lime arrow. Increasing ocean depth represented by a darkening blue scale. Base map made using the ggOceanMaps R package (Vihtakari 2022) with data from Natural Earth Data, Amante & Eakins (2009), Geonorge.no, and the General Bathymetric chart of the Oceans, and Inkscape 1.1.

## Methods

### Sample collection

Samples used were those originally described in Jacobsen & Hansen (2001). Samples were collected as part of an experimental longline fishery that occurred in two seasons, referred to as the 1993 season and the 1994 season. Each season lasted from November of the preceding year until March of the following year, with no fishing in January due to poor sea conditions. To illustrate, the 1993 season saw sampling occur in November and December of 1992 and in February and March of 1993. Fish were caught on longlines baited with European sprat, *Sprattus sprattus*, with lines left to soak from around dawn to noon. Due to its experimental nature, the fishery was exempt from the minimum hook size commonly imposed on longline fisheries (KH, pers. obs.). Thus, the distribution of fork lengths and, therefore, sea ages, observed amongst sampled fish should be, *ceteris paribus*, unbiased by hook size selectivity and representative of the true sea age structure of the fish at the time of sampling. However, size selectivity of the sampled fish might still have occurred if bait size favoured larger conspecifics (Ingólfsson et al. 2017; see Discussion). The date and capture location of each sampled fish were recorded. Scales were sampled and the sea ages of individual fish determined by the number of winters they had spent at sea. In this study, individuals were designated as either one-sea winter or multi-sea winter (referred to as 1SW and MSW, respectively, from hereafter). This combination of date- and location-specific catch records, coupled with sea age phenotypes, allowed for the proportion of MSW salmon present in the fishery to be estimated. See Jacobsen & Hansen (2001) for further details of the fishery and sample processing.

### Genotyping and genetic stock identification

The genetic baseline used to assign fish to specific reporting groups was generated by combining SNP data from previously published Atlantic salmon studies (Table S1) with unpublished data for the Alta and Teno rivers. Where DNA had been pooled (*n* = 40 populations), individual genotypes were generated from the allele frequency estimates, assuming both Hardy-Weinberg and linkage equilibrium (*n* = 40 per population). This was done using a bespoke software as genetic stock identification, GSI, requires individual genotype data rather than allele frequency estimates (see details in Ozerov et al. 2013). Given that individual populations in the baseline were geographically dispersed and we were interested in broad-scale macroecological patterns, we used agglomerative reporting groups instead of specific populations to define the origin of the salmon sampled from around the Faroes. Reporting groups (Fig. 1) were defined using a neighbour joining tree (Fig. S2), information from previous studies exploring population-level genetic variation in Atlantic salmon (e.g. Ozerov et al. 2013; Ozerov et al. 2017; Gilbey et al. 2017), as well as expert opinion from this study’s authors. DNA from Faroese-sampled and Teno salmon was extracted from archived scales. For Alta fish, fin clips stored at −20°C were used in lieu of scales. Extraction was conducted using a QIAamp 96 DNA QIAcube HT Kit (Qiagen) following the manufacturer’s protocol. Samples were further genotyped by targeted sequencing at 167 SNP markers using a GTSeq approach (Campbell et al. 2015) as outlined in Aykanat et al. (2020).

The origin for each successfully genotyped individual was estimated using the conditional maximum likelihood GSI methodology (Millar, 1987) implemented in ONCOR (Kalinowski et al. 2008). Since estimation of population origin using GSI methods can be affected by the population composition of the mixture sample (Pella & Masuda, 2001), we divided the total number of fish to be assigned into the individual fishing seasons (1993, 1994), seasons (autumn, winter), and location (north, south) in which they were caught. The inclusion of such spatio-temporal variation in stock distribution can improve GSI sensitivity (Vähä et al. 2017 as read in Svenning et al. 2019). In total, 1674 samples were divided into six temporally and spatially distinct subsets for the GSI analysis. The assignment probability (p) threshold for reporting groups was set at ≥ 0.7 (Vähä et al. 2011, 2014; Bradbury et al. 2015).

The ability of our baseline to successfully assign fish to their reporting group of origin was evaluated with 100% simulation and leave-one-out cross-validation methods (Anderson et al. 2008) implemented in ONCOR (Kalinowski et al. 2008). For the 100% simulation evaluation, we set the mixture sample sizes to 200 and simulations were repeated 100 times for both the individual populations as well as for the defined reporting groups. The leave-one-out test was performed by removing fish from baseline populations (one at a time) and then estimating their origin to evaluate how well a fish could be assigned to their population or reporting group of origin. Fish with incomplete genotypes (i.e. no value at one or more loci) were not used during the leave-one-out cross-validation, however, they were retained in the baseline in order to estimate the origin of fish (Kalinowski et al. 2008).

While our GSI baseline contained samples from at least one population in each of the major Atlantic salmon phylogenetic groups (Bourret et al. 2013), downstream analyses focused on six eastern Atlantic reporting groups (Ireland and UK, Southern Norway, Northern Norway, Eastern Finnmark/North Kola, Teno, and Eastern Barents Sea/White Sea), as well as the North American reporting group. This was due to no fish being assigned to Icelandic, Baltic, or Bay of Biscay reporting groups (Table S1).

### Estimation of MSW proportion among Faroese-sampled fish and among the deep-sea pre-fisheries abundance

#### (1) Faroese fish

We estimated the proportion, and associated uncertainty, of MSW fish sampled around the Faroes from each of the reporting groups using a generalized linear model with a logit link and a binomial response variable (success = MSW, failure = 1SW), reporting group as a fixed factor, a categorical link function, and a flat prior on the probability scale. For the fixed effects, we used the prior *N*(0, I × (1 + *π*^2^/3)), where 0 is a vector of zeros and I is the identity matrix, both with dimensions equal to the number of reporting groups (six in this case). We fixed the residual variance at one. The model was run using MCMCglmm (Hadfield 2010), which allowed for the uncertainty around each proportion estimate to itself be estimated (since parameter values estimated using MCMCglmm are probabilistic due to the software package implementing a Bayesian paradigm). The median of the generated posterior distribution and the 95% credible intervals (95% CI) were used to express the MSW proportion and its associated uncertainty, respectively. Chains were run for 2,000,000 iterations with the first 500,000 iterations discarded as burn-in. Realisations of the Markov chain were sampled every 150 iterations.

#### (2) Pre-fisheries abundance

The natural abundance of salmon in the North Atlantic Ocean prior to Faroese and coastal fisheries is termed the pre-fishery abundance (PFA). We used the run-reconstruction model (Potter et al. 2004), described in the stock annex of the ICES Working Group on North Atlantic Salmon (WGNAS; ICES, 2021) Briefly, this is done by first correcting abundance in the oceans upwards, using the reported catch of 1SW and MSW salmon of each stock, to estimate the number of salmon returning to the coastal ‘homewaters’ of stocks by taking into account the non-reported catch and an exploitation rate (specified as uniform distributions). These numbers were further corrected (increased) by accounting for natural mortality (assumed to be 2-4% per month, uniformly distributed) across the average number of months until the return of both age classes in the different stocks. This number is now an estimation of the PFA as of 1^st^ January in a given year. The model uses Monte Carlo simulation (9999 iterations) to quantify PFA with uncertainty. We used the data collated by the WGNAS (ICES, 2021) to quantify 1SW and MSW PFA for the years 1993 and 1994. We did this separately for Ireland and UK (ICES WGNAS regions: England & Wales, Scotland east and west, Northern Ireland Foil Fisheries Area and DAERA area, and Ireland), South Norway (Norway south-east, south-west, and middle), Northern Norway (Norway north), Teno (Finland), Eastern Finnmark/North Kola (Russia Kola Peninsula, Barents Sea Basin), and Eastern Barents Sea/White Sea (Russia Kola Peninsula, White Sea Basin, Archangelsk and Karelia, and Pechora River). The reconstructed abundance data was then used to estimate the proportion of MSW fish in the deep seas. We used the median and 95% CI to describe the MSW proportion and its uncertainty for each of the reporting groups.

To further explore the observed differences in MSW proportions (see Results), we estimated the relative likelihood of occurrence of a given sea age class for each reporting group sampled around the Faroes by normalising the number of salmon assigned to each reporting group/age class to the PFA of each reporting group. This was done as follows;

1. Relative likelihood of occurrence 1SW_*j*_ = N_FO*i*_ × (1 - P_MSW*i*_) / PFA_*i,j*_;
2. Relative likelihood of occurrence MSW_*j*_ = N_FO*i*_ × P_MSW*i*_ / PFA_*i,j*_

where subscripts *j* and *i* denote the regional group and sea age group of fish, respectively, N_FO_ is number of fish assigned to each regional group, P_MSW_ is the posterior distribution of proportion of MSW fish, and PFA is the Monte Carlo sampling distribution of the pre-fishery abundance. Note, relative likelihood of occurrence is an arbitrary measure that provides a convenient means to compare between sea ages and reporting groups the likelihood of a salmon migrating to the Faroes, relative to their abundance at sea.

#### (3) Consistency of pattern across MSW age classes

Is increasing sea age associated with increasing likelihood of presence around the Faroes or does the pattern exist solely as a dichotomy between 1SW and MSW fish? To further explore this, we decomposed the MSW component of the Northern and Southern Norway reporting groups into their constituent 2SW and 3SW components. We used the same run-reconstruction model as above but with an additional age class (3SW). This model used the MSW exploitation rate and unreported catch reported to ICES for both 2SW and 3SW fish. The necessary 2SW/3SW catch data is given in Table S4.

We then calculated the proportion of 3SW fish to 2SW fish around the Faroes separately for both Southern and Northern Norway. We then repeated this calculation, but instead using the PFA for each age class. Finally, we divided the 3SW to 2SW proportion for the Faroes by the 3SW to 2SW proportion for the PFA. This figure provides an estimate of the proportional increase of 3SW salmon relative to 2SW salmon around the Faroes, normalised to the PFA of each age class.

Analyses and plots can be recreated using the code and data stored at https://github.com/Helsinki-Ronan/MSW-Faroes-migration-paper. All analyses were conducted in R version 4.1.2 (R Core Team 2021). Visual inspection of Markov chains suggested that models mixed well and that model parameter estimates had converged upon stable distributions.

## Results

### Genetic stock identification

1616 fish were confidently assigned back to their reporting group and used in further analyses. A total of 454 fish (28.1%) assigned back to Ireland and UK, 817 (50.6%) to Southern Norway, 115 (7.1%) to Northern Norway, 24 (1.5%) to Teno, 10 (0.6%) to Eastern Finnmark, North Kola, 110 (6.8%) to the Eastern Barents, White Sea (Table 2), and 86 (5.3%) to North America. See Tables S2 and S3 for assignment confidence at the population and reporting group levels, respectively.

### Estimation of MSW proportions

The MSW proportion for each reporting group found around the Faroes was consistently larger than the proportion of MSW individuals among PFA for each of the six aforementioned European reporting groups (Fig. 2a; Table 1). For individual reporting groups, the proportion of MSW Atlantic salmon sampled from around the Faroes ranged from 0.749 (95% CI: 0.702, 0.793) for Ireland and UK to 0.995 for Northern Norway (95% CI: 0.978, 0.999). The proportion of MSW for Ireland and UK was strikingly lower than in all other reporting groups (with pairwise non-overlapping 95% credible intervals) except when compared to the Eastern Finnmark, North Kola group which had a high observed proportion but this was likely due to high uncertainty of the estimate as a result of low sample size (Fig. 2a; Table 1).

**Figure 2:**
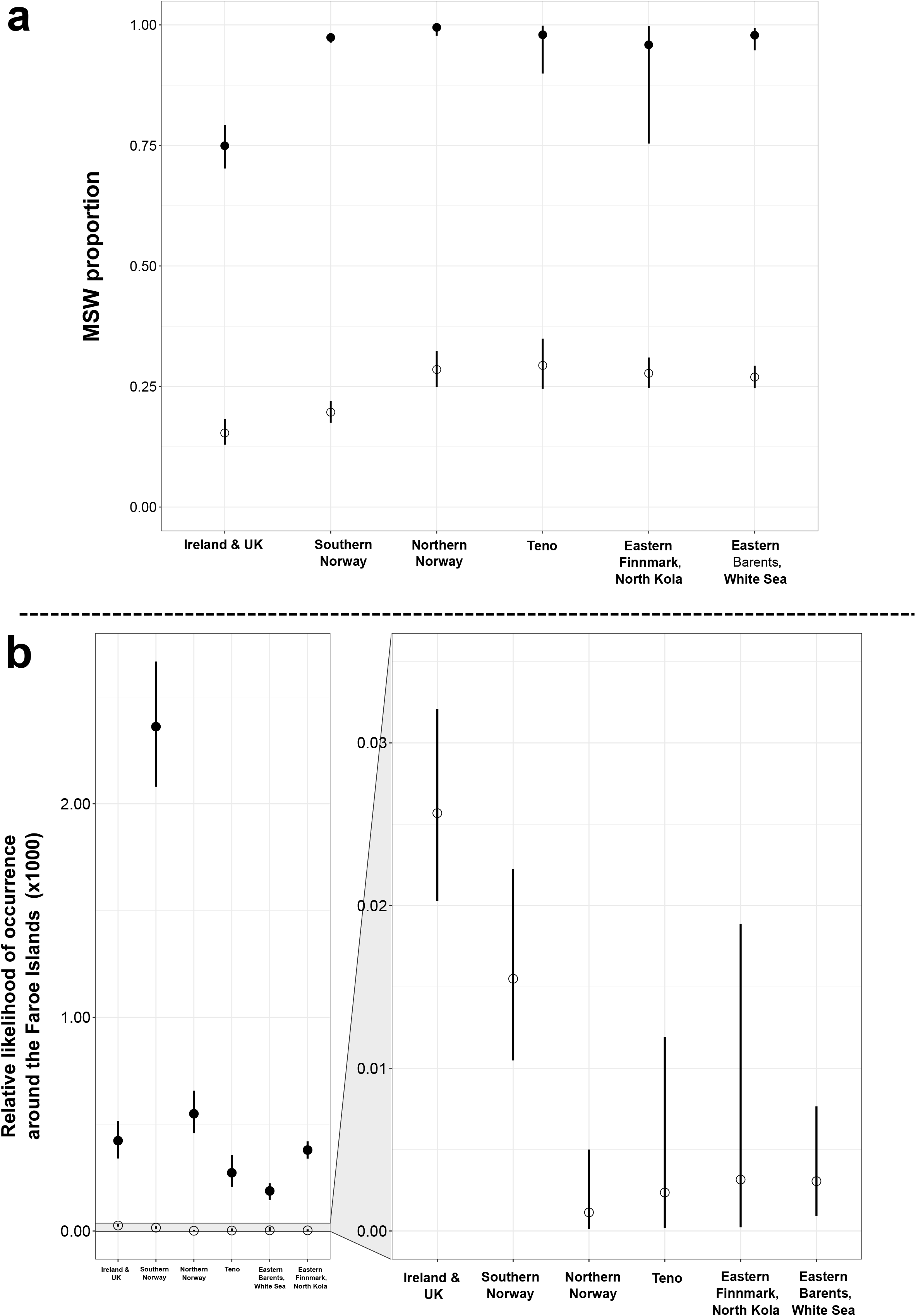
(a) Differences in the proportion of multi-sea winter (MSW) Atlantic salmon sampled from around the Faroe Islands (filled circles) during the 1993 and 1994 fishing seasons and the proportion of MSW fish among the estimated pre-fisheries abundance, totaled across the years 1993-1994 (open circles); (b) Differences in the relative likelihood of occurrence of MSW (filled circles) and one-sea winter 1SW (open circles) Atlantic salmon with respect to their agespecific, deep-sea pre-fisheries abundance. For both (a) and (b), point estimates and uncertainty represented with medians and 95% credible intervals, respectively.

**Table 1:** The numbers (nMSW) and proportion of multi-sea winter (MSW) Atlantic salmon assigning to each of the six eastern Atlantic reporting groups sampled in an experimental fishery (Faroe Islands), and the proportion of MSW fish among the deep-sea pre-fisheries abundance (PFA) estimated for each reporting group. The number of one-sea winter fish used to calculate the Faroese MSW proportion are denoted by ‘n1SW’.

| Reporting Group | Location | n1SW, nMSW | MSW proportion | 2.5% credible interval | 97.5% credible interval |
| --- | --- | --- | --- | --- | --- |
| Ireland and UK | Faroe Islands | 130, 324 | 0.749 | 0.702 | 0.793 |
|  | PFA |  | 0.154 | 0.129 | 0.183 |
| Southern Norway | Faroe Islands | 32, 785 | 0.974 | 0.963 | 0.982 |
|  | PFA |  | 0.197 | 0.175 | 0.220 |
| Northern Norway | Faroe Islands | 0, 115 | 0.995 | 0.978 | 0.999 |
|  | PFA |  | 0.285 | 0.249 | 0.324 |
| Teno | Faroe Islands | 0, 24 | 0.980 | 0.899 | 0.998 |
|  | PFA |  | 0.294 | 0.245 | 0.349 |
| Eastern Finnmark, North Kola | Faroe Islands | 0, 10 | 0.959 | 0.740 | 0.997 |
|  | PFA |  | 0.277 | 0.247 | 0.310 |
| Eastern Barents, White Sea | Faroe Islands | 3, 107 | 0.979 | 0.947 | 0.993 |
|  | PFA |  | 0.270 | 0.247 | 0.293 |

The relative likelihood of occurrence for 1SW salmon around the Faroes relative to 1SW PFA was higher for Ireland and UK (0.026, 95% CI: 0.020, 0.032) as well as the Southern Norway reporting groups (0.016, 95% CI: 0.010, 0.022) compared to the other four reporting groups where the relative likelihood of occurrence was virtually zero (Fig. 2b). The relative likelihood of occurrence for MSW salmon around the Faroes relative to their PFA was highest for Southern Norway (2.36, 95% CI: 2.08 2.67), with Northern Norway displaying the second highest rate of occurrence (0.55, 95% CI: 0.46, 0.66). Strikingly, MSW salmon from Ireland and UK were as likely to be found around the Faroes as MSW fish from more north-eastern reporting groups (Teno/Eastern Barents, White Sea), despite Ireland and UK being far closer geographically to the Faroes than these latter groups (Fig. 1).

### Consistency of pattern

For Southern Norway, 3SW Atlantic salmon were 8.31 (95% CI: 6.90, 9.98;) times more likely to be present around the Faroes than 2SW fish, with respect to the PFA of each sea age. For Northern Norway, this figure was 1.76 (95% CI: 1.11, 2.77; Fig. 3).

**Figure 3:**
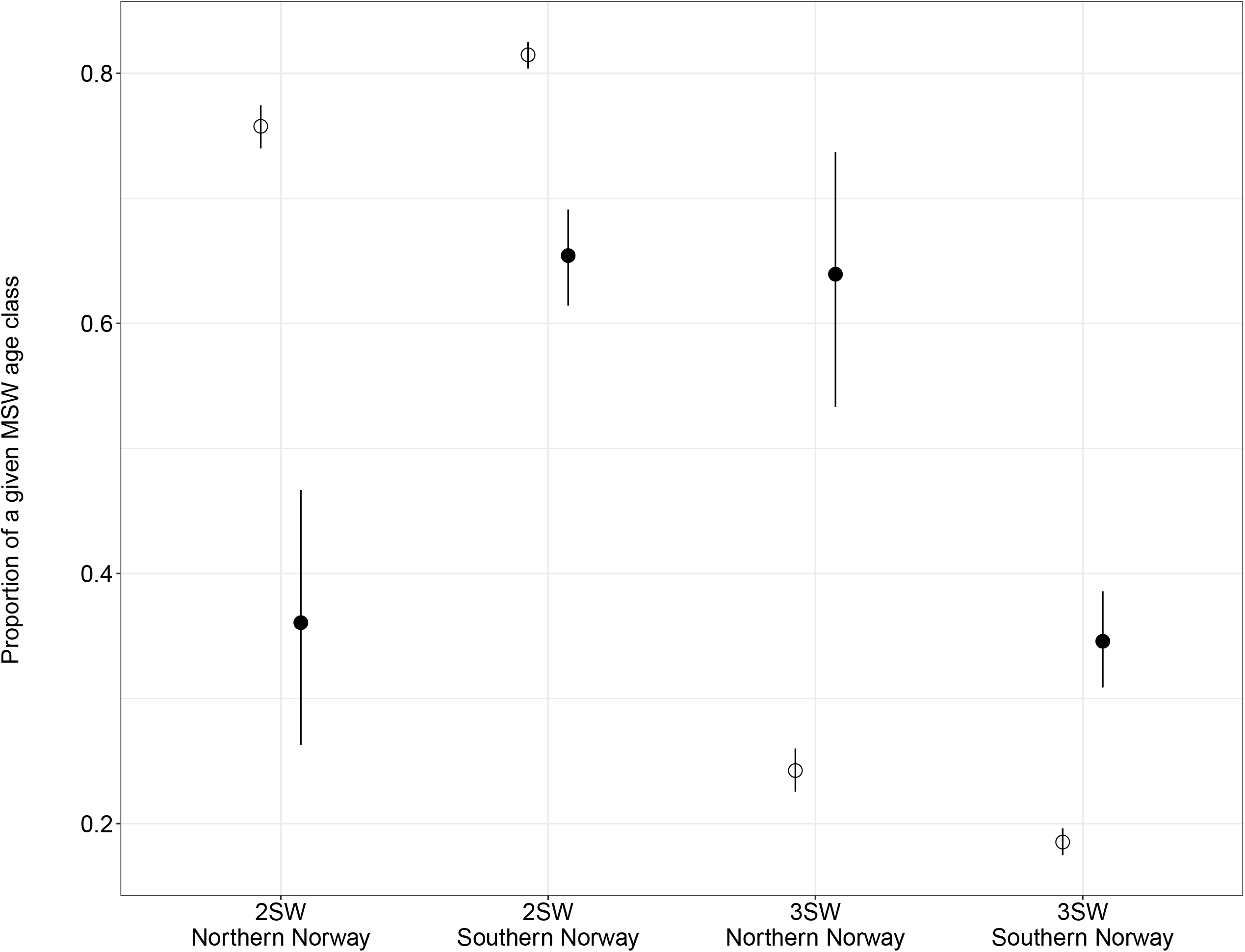
Proportion of 2SW and 3SW Atlantic salmon within the multi-sea winter (MSW) age class sampled around the Faroe Islands (filled circles) and estimated for the deep-sea prefisheries abundance (open circles) for the Southern Norway and Northern Norway reporting groups. Point estimates and uncertainty represented with medians and 95% credible intervals, respectively.

## Discussion

### Faroese feeding ground versus pre-fisheries abundance MSW proportions

By genotyping spatially- and temporally-explicit Atlantic salmon individuals of known sea age sampled around the Faroe Islands, we were able to demonstrate that MSW fish from six European reporting groups use this oceanic feeding ground in greater proportions than 1SW salmon from the same groups (Fig. 2a). In particular, MSW fish and especially those from more northern areas appear to make greater use of multiple, large-scale ecosystems (Barents Sea, Norwegian Sea) over the course of their marine migration than do 1SW fish from those same stocks, at least during the sampled study period. Such phenotype-specific exploitation of spatially dislocated areas suggests that Atlantic salmon might preferentially migrate to specific feeding grounds and that fish spending more than one winter at sea might move between different feeding areas. The greater-than-expected number of 3SW fish around the Faroes provides further evidence that older fish preferentially migrate there (Fig. 3).

Intriguingly, MSW salmon from the Ireland and UK reporting group were only as likely to be found around the Faroes as MSW fish from much more geographically distant north-eastern reporting groups (Fig. 2b). In turn, 1SW fish from Ireland and UK were generally more likely to be present at the Faroes than 1SW fish from the other reporting groups, with Southern Norway being an exception to this general pattern. These two groups displayed similar rates of migration to the Faroes for 1SW fish (Fig. 2b), while MSW fish from Southern Norway were more likely to migrate to the Faroes than MSW fish from the rest of the reporting groups. These results suggest that, despite their proximity to the Faroes, many MSW salmon from Ireland and UK do not migrate there. The main feeding ground of these MSW salmon from Ireland and UK that don’t utilise the Faroes remains unknown. One potential candidate is the Labrador Sea to the west of Greenland (Fig. 1). Bradbury et al. (2016) found that 22% of salmon sampled from the Labrador Sea displayed European heritage. Subsequent fine-scale GSI has assigned many salmon sampled in the Labrador Sea back to Ireland and UK (Bradbury et al. 2021). Furthermore, a large number of tagged salmon from Scotland, England, and Wales have been recovered in the Labrador Sea, providing further support for an Ireland and UK-Labrador Sea migration route (Swain, 1980).

How this general pattern of spatial resource use emerges appears to be related to the flow of currents around the North Atlantic. Based on a combination of tagging data, isotope analysis, and fishery patterns, Dadswell et al. (2010) suggested Atlantic salmon oceanic migrations are determined by the North Atlantic subpolar gyre. This gyre flows counter-clockwise between North America and north-western Europe with various regional currents and eddies within it (Fig. 1). Many salmon post-smolts from Ireland and UK likely take advantage of these regional oceanographic features and migrate directly to the Faroes, with the eddy system between the islands and the Vøring Plateau likely helping them to stay in this general area (Fig. 1). This supposition is supported by an annual concentration of southern origin post-smolts at the Vøring Plateau (Gilbey et al. 2021). The lower-than-expected proportion of MSW salmon from Ireland and UK around the Faroes, along with the results of Bradbury et al. (2016, 2021) and Swain (1980), suggests that a portion of the Ireland and UK post-smolts in this concentration of fish might undergo a further migration to the Labrador Sea.

The feeding grounds of Atlantic salmon from the north-eastern reporting groups are not as well-known as the feeding grounds of more southernly groups (Strøm et al. 2018; Rikardsen & Dempson, 2011; Rikardsen et al. 2021). However, most of our knowledge of these feeding grounds comes from tagged adult salmon who had previously spawned at least once and whose oceanic migration behaviour may differ to maiden fish (see also Rikardsen et al. 2021). It is probable that fish from these north-eastern reporting groups are less likely to undergo long oceanic migrations, such as to the Faroes, and remain on feeding grounds closer to their source populations within the Barents and White Seas (Jensen et al. 1999; Rikardsen et al. 2008). Gilbey et al. (2021) found no post-smolts from the north-eastern reporting groups in the Faroes which they suggested was likely due to such fish being carried in the opposite direction by eastward currents (Fig. 1). Despite this potential initial displacement, we assigned a substantial number of MSW fish back to the north-eastern reporting groups (*n* = 257; Table 1). The relative likelihood of occurrence of MSW salmon around the Faroes from these groups was sometimes as high as that of Ireland and UK (Fig. 2b), suggesting this is a significant feeding ground for north-eastern MSW fish.

### North American, Icelandic, and Bay of Biscay salmon

Eighty-six individuals were assigned to salmon populations in North America (5.3% of assigned fish). The total pre-fisheries abundance for North America during the study period (1993-1994) was 1,103,500 fish (ICES, 2021) compared to 9,213,265 fish from the six European reporting groups in this study to which salmon were successfully assigned. All else being equal, this means that Atlantic salmon from North America were 2.2 times less likely to be found around the Faroes relative to fish from Europe. Overlap of Atlantic salmon from North America and from Europe in the ocean is a known phenomenon (Hansen & Jacobsen, 2003; Dadswell et al. 2010; Bradbury et al. 2016, 2021). Spares et al. (2007) used isotope analysis to infer that 14.2% of 141 Canadian salmon had fed east of the Faroes. Additionally, fish tagged in the Faroes have been controlled in North American populations (Dadswell et al. 2010). Similar to Bradbury et al. (2016, 2021) and Gilbey et al. (2017), this work corroborates these previous studies by providing direct genetic evidence for such movements by salmon. The high percentage 89.5% (*n* = 77 out of 86) of MSW individuals among North American-assigned fish from this study tentatively suggests that the feeding grounds around the Faroes are indeed sought out by MSW salmon from North American stocks.

In this study, no fish were assigned to Iceland despite its proximity to the Faroes. Gilbey et al. (2021) found a lower-than-expected number of Icelandic post-smolts in the same geographic area as this study. Similarly, only one salmon out of 87 recaptures from an ocean tagging programme conducted around the Faroes was controlled in Iceland (Hansen & Jacobsen, 2003). Using estimated marine distributions, Guðjónsson et al. (2015) suggest that Icelandic fish are more likely to migrate west of Iceland towards the Irminger Sea. Similarly, two post-spawned Atlantic salmon tagged in Iceland migrated directly to this area (Rikardsen et al. 2021). Why the majority of Icelandic fish go westward instead of migrating east to the Faroes remains yet to be explored. Similarly, none of the 1616 Atlantic salmon were assigned back to the Bay of Biscay reporting group. This was also surprising given Biscay salmon are believed to share a northward migration route with fish from Ireland and UK (Otero et al. 2014). There are only two tag recoveries of Biscay salmon in the Faroese tag database (Ó’Maoiléidigh et al. 2018). Our results suggest that Biscay fish, regardless of sea age, do not feed in the ocean around the islands during the November-March period of the year. While it is expected that fish from this reporting group would be present in low numbers around the Faroes (based on post-smolt distribution – Gilbey et al. 2021), this raises the question of where do these post-smolts subsequently migrate to (though one out of 14 tagged post-spawned adult salmon from the Lérez river in Spain is known to have migrated to the west Irminger Sea - Rikardsen et al. 2021). Their absence from data sets might also reflect the generally poor status of Biscay salmon populations, with many already extirpated or declining (Parrish et al. 1998).

### Evolutionary and ecological implications in a changing ocean

Changes in the spatial availability of prey resources during the marine stage of Atlantic salmon might be changing the costs and benefits of undergoing migrations, such as that to the Faroes. Vollset et al. (2022) documented lower abundances of key prey items taken by salmon during the early marine stage in the Norwegian Sea from 2005 onwards. This decrease in resources (e.g. a reduction in zooplankton) coincided with a decrease in both the abundance and body condition of 1SW salmon (Utne et al. 2021) which could be the result of evolution towards an older age structure as a response to many fish not reaching their maturity threshold in their first year at sea. Interestingly, Czorlich et al. (2022) demonstrated how a reduction in capelin availability was driving evolution of age structure towards early maturation among Teno salmon. While exhibiting opposite responses to reduced prey availability, the previous two studies share a common theme of changes in age structure resulting from altered availability of marine food resources. If our hypothesis is correct and different age classes and different stocks display variation in their utilization of the oceans around the Faroes then any deleterious changes in the marine conditions around the Faroes will have age- and stock-specific responses. As discussed, Vollset et al. (2022) document a contemporary regime shift and its negative impact on the abundance of Atlantic salmon prey items. Other such regime shifts are predicted to occur in the future (Beaugrand et al. 2008), each of which carries the potential of adverse effects on the Faroese marine food web as well as other oceanic ecosystems. Future research should aim to quantitatively assess the likelihood of such regime shifts affecting food webs and, thus, the vulnerability of different age classes across individual salmon stocks.

### Potential limitations

Like many studies using data from opportunely collected samples of wild individuals, any inference of our results must be done so with knowledge of the data’s shortcomings. As discussed in the Methods, the experimental longline fishery was not limited by a minimum hook size, thus, precluding hook size selectivity from biasing our data. The size of bait used on the hooks could potentially introduce bias by preferentially targeting larger fish (Ingólfsson et al. 2017). However, we do not believe this bias qualitatively impacts our results given that post-smolt/one-sea winter salmon during our sampling period were large enough (mean fork length ± one standard deviation = 48.79 ± 4.33 cm) so as not to be gape-limited with respect to sprat (average size sprat = 10.1 cm – FishBase). Also, the results from splitting the Southern and Norwegian groups’ MSW components into the 2SW and 3SW age classes (among which the size of fish entirely precludes gape-limitation) demonstrates that the oceans around the Faroes are indeed enriched with 3SW salmon compared to 2SW fish. Taken together, these results show that there was no bait bias between these two age classes. This reinforces the supposition that the ocean around the Faroes is exploited more by older age classes and that the observed proportion of 3SW salmon among our samples is representative of the true proportion of this age class (Fig. 3).

As the fish in this study were all sampled in late autumn and early winter, and the fishery was almost exclusively located to the north and west of the Faroes, the data are both temporally and spatially unbalanced. However, Ó’Maoiléidigh et al. (2018) demonstrated that salmon caught around the Faroes during May-June were also predominantly MSW fish. This suggests that the large proportion of MSW salmon sampled by the experimental fishery is observed at other times of the year and, thus, supports the idea that the seas around the Faroes predominantly support MSW fish. The effect of spatial unbalance on how representative our results are is more difficult to determine due to a lack of data to bring to bear on the question.

## Conclusion

Using a combination of genetic stock assignment and run-reconstruction models, we demonstrated that the proportion of MSW Atlantic salmon sampled on their feeding grounds around the Faroe Islands was consistently higher than the estimated MSW proportion among deep-seas pre-fisheries abundance. This pattern was observed across six eastern Atlantic reporting groups. Our results suggest that the seas around the Faroes are a preferred feeding ground for MSW salmon. Preferentially migrating to an area in order to exploit its resources is a behaviour likely mediated, in part, by some evolved feeding strategy. If the MSW proportion of a given reporting group is more likely to exploit a specific spatial resource (such as the Faroese feeding ground) than 1SW fish from the same reporting group, then changes in the spatial and temporal availability of such resources are likely to differentially effect intra-group age classes. Similarly, if MSW individuals from different reporting groups utilise the ocean around the Faroes to greater or lesser extents, then changes in feeding conditions at the Faroes will also induce reporting group-specific changes to the MSW component of reporting groups. Understanding the temporal and spatial dynamics of oceanic resource use both within (i.e. age classes) and between stocks is crucial in making anticipatory predictions as to how different Atlantic salmon populations might be affected by changes to marine food webs.

## Acknowledgements

The authors would like to thank the crew of the M/S Hvítiklettur, Nina Suomalainen, University of Helsinki DNA Sequencing and Genomics Lab, and all others who have been involved in the collection and processing of the samples used in this study. This study was funded by Academy of Finland grants to TA (project numbers: 1328860 and 1325964). RJOS was supported by an Ella and Georg Ehrnrooth Postdoctoral Researcher Grant. Salary to GHB and KH was funded by the Norwegian Research Council (projects 275862 and 280308).

## References

1. Aas, Ø., Policansky, D., Einum, S., & Skurdal, J. (2010). Salmon Ecological Research and Conservation. In Ø. Aas, S. Einum, A. Klemetsen, & J. Skurdal (Eds.), Atlantic Salmon Ecology (pp. 445–456). Wiley-Blackwell. https://doi.org/10.1002/9781444327755.ch17

2. Amante, C. & Eakins B.W. (2009). ETOPO1 1 Arc-Minute Global Relief Model: Procedures, Data Sources and Analysis. NOAA Technical Memorandum NESDIS NGDC-24. National Geophysical Data Center, NOAA. Distributed under the U.S. Government Work license.

3. Anderson, E. C., Waples, R. S., & Kalinowski, S. T. (2008). An improved method for predicting the accuracy of genetic stock identification. Canadian Journal of Fisheries and Aquatic Sciences, 65(7), 1475–1486. https://doi.org/10.1139/F08-049

4. Armstrong, M. P., Dean, M. J., Hoffman, W. S., Zemeckis, D. R., Nies, T. A., Pierce, D. E., Diodati, P. J., & McKiernan, D. J. (2013). The application of small scale fishery closures to protect Atlantic cod spawning aggregations in the inshore Gulf of Maine. Fisheries Research, 141, 62–69. https://doi.org/10.1016/j.fishres.2012.09.009

5. Aykanat, T., Rasmussen, M., Ozerov, M., Niemelä, E., Paulin, L., Vähä, J., Hindar, K., Wennevik, V., Pedersen, T., Svenning, M., & Primmer, C. R. (2020). Life-history genomic regions explain differences in Atlantic salmon marine diet specialization. Journal of Animal Ecology, 89(11), 2677–2691. https://doi.org/10.1111/1365-2656.13324

6. Ayllón, D., Nicola, G. G., Elvira, B., & Almodóvar, A. (2019). Optimal harvest regulations under conflicting tradeoffs between conservation and recreational fishery objectives. Fisheries Research, 216, 47–58. https://doi.org/10.1016/j.fishres.2019.03.021

7. Ayllón, D., Railsback, S. F., Almodóvar, A., Nicola, G. G., Vincenzi, S., Elvira, B., & Grimm, V. (2018). Eco-evolutionary responses to recreational fishing under different harvest regulations. Ecology and Evolution, 8(19), 9600–9613. https://doi.org/10.1002/ece3.4270

8. Beaugrand, G., Edwards, M., Brander, K., Luczak, C., & Ibanez, F. (2008).: Causes and projections of abrupt climate-driven ecosystem shifts. Ecology Letters, 11(11), 1157–1168. https://doi.org/10.1111/j.1461-0248.2008.01218.x

9. Bourret, V., Kent, M. P., Primmer, C. R., Vasemägi, A., Karlsson, S., Hindar, K., McGinnity, P., Verspoor, E., Bernatchez, L., & Lien, S. (2013). SNP-array reveals genome-wide patterns of geographical and potential adaptive divergence across the natural range of Atlantic salmon (*Salmo salar*). Molecular Ecology, 22(3), 532–551. https://doi.org/10.1111/mec.12003

10. Bradbury, I. R., Hamilton, L. C., Rafferty, S., Meerburg, D., Poole, R., Dempson, J. B., Robertson, M. J., Reddin, D. G., Bourret, V., Dionne, M., Chaput, G., Sheehan, T. F., King, T. L., Candy, J. R., & Bernatchez, L. (2015). Genetic evidence of local exploitation of Atlantic salmon in a coastal subsistence fishery in the Northwest Atlantic. Canadian Journal of Fisheries and Aquatic Sciences, 72(1), 83–95. https://doi.org/10.1139/cjfas-2014-0058

11. Bradbury, I. R., Hamilton, L. C., Sheehan, T. F., Chaput, G., Robertson, M. J., Dempson, J. B., Reddin, D., Morris, V., King, T., & Bernatchez, L. (2016). Genetic mixed-stock analysis disentangles spatial and temporal variation in composition of the West Greenland Atlantic Salmon fishery. ICES Journal of Marine Science, 73(9), 2311–2321. https://doi.org/10.1093/icesjms/fsw072

12. Bradbury, I. R., Lehnert, S. J., Messmer, A., Duffy, S. J., Verspoor, E., Kess, T., Gilbey, J., Wennevik, V., Robertson, M., Chaput, G., Sheehan, T., Bentzen, P., Dempson, J. B., & Reddin, D. (2021). Range-wide genetic assignment confirms long-distance oceanic migration in Atlantic salmon over half a century. ICES Journal of Marine Science, 78(4), 1434–1443. https://doi.org/10.1093/icesjms/fsaa152

13. Campbell, N. R., Harmon, S. A., & Narum, S. R. (2015). Genotyping-in-Thousands by sequencing (GT-seq): A cost effective SNP genotyping method based on custom amplicon sequencing. Molecular Ecology Resources, 15(4), 855–867. https://doi.org/10.1111/1755-0998.12357

14. Czorlich, Y., Aykanat, T., Erkinaro, J., Orell, P., & Primmer, C. R. (2022). Rapid evolution in salmon life history induced by direct and indirect effects of fishing. Science, 376(6591), 420–423. https://doi.org/10.1126/science.abg5980

15. Dadswell, M. J., Spares, A. D., Reader, J. M., & Stokesbury, M. J. W. (2010). The North Atlantic subpolar gyre and the marine migration of Atlantic salmon Salmo salar: The ‘Merry-Go-Round’ hypothesis. Journal of Fish Biology, 77(3), 435–467. https://doi.org/10.1111/j.1095-8649.2010.02673.x

16. Eikeset, A. M., Richter, A., Dunlop, E. S., Dieckmann, U., & Stenseth, N. Chr. (2013). Economic repercussions of fisheries-induced evolution. Proceedings of the National Academy of Sciences, 110(30), 12259–12264. https://doi.org/10.1073/pnas.1212593110

17. Fleming, I. A. (1998). Pattern and variability in the breeding system of Atlantic salmon (*Salmo salar*), with comparisons to other salmonids. Canadian Journal of Fisheries and Aquatic Sciences, 55(S1), 59–76. https://doi.org/10.1139/d98-009

18. FishBase (2022). Sprattus sprattus summary page. Available at: <https://www.fishbase.se/summary/Sprattus-sprattus.html> [Accessed 1 July 2022].

19. Gilbey, J., Utne, K. R., Wennevik, V., Beck, A. C., Kausrud, K., Hindar, K., Garcia de Leaniz, C., Cherbonnel, C., Coughlan, J., Cross, T. F., Dillane, E., Ensing, D., García-Vázquez, E., Hole, L. R., Holm, M., Holst, J. C., Jacobsen, J. A., Jensen, A. J., Karlsson, S.,… Verspoor, E. (2021). The early marine distribution of Atlantic salmon in the North-east Atlantic: A genetically informed stock-specific synthesis. Fish and Fisheries, 22(6), 1274–1306. https://doi.org/10.1111/faf.12587

20. Gilbey, J., Wennevik, V., Bradbury, I. R., Fiske, P., Hansen, L. P., Jacobsen, J. A., & Potter, T. (2017). Genetic stock identification of Atlantic salmon caught in the Faroese fishery. Fisheries Research, 187, 110–119. https://doi.org/10.1016/j.fishres.2016.11.020

21. Gross, M. R., Coleman, R. M., & McDowall, R. M. (1988). Aquatic productivity and the evolution of diadromous fish migration. Science, 239(4845), 1291–1293. https://doi.org/10.1126/science.239.4845.1291

22. Guðjónsson, S., Einarsson, S. M., Jónsson, I. R., & Guðbrandsson, J. (2015). Marine feeding areas and vertical movements of Atlantic salmon (*Salmo salar*) as inferred from recoveries of data storage tags. Canadian Journal of Fisheries and Aquatic Sciences, 72(7), 1087–1098. https://doi.org/10.1139/cifas-2014-0562

23. Hadfield, J. D. (2010). MCMC methods for multi-response generalized linear mixed models: The MCMCglmm R package. Journal of Statistical Software, 33(2), 1–22. https://doi.org/10.18637/jss.v033.i02

24. Hansen, L. P., & Jacobsen, J. A. (2003). Origin and migration of wild and escaped farmed Atlantic salmon, Salmo salar L., in oceanic areas north of the Faroe Islands. ICES Journal of Marine Science, 60(1), 110–119. https://doi.org/10.1006/jmsc.2002.1324

25. Harada, A. E., Lindgren, E. A., Hermsmeier, M. C., Rogowski, P. A., Terrill, E., & Burton, R. S. (2015). Monitoring spawning activity in a Southern California Marine Protected Area using molecular identification of fish eggs. PLOS ONE, 10(8), e0134647. https://doi.org/10.1371/journal.pone.0134647

26. Hawley, K. L., Rosten, C. M., Christensen, G., & Lucas, M. C. (2016). Fine-scale behavioural differences distinguish resource use by ecomorphs in a closed ecosystem. Scientific Reports, 6(1), 24369. https://doi.org/10.1038/srep24369

27. Hernández, C. M., Witting, J., Willis, C., Thorrold, S. R., Llopiz, J. K., & Roţjan, R. D. (2019). Evidence and patterns of tuna spawning inside a large no-take Marine Protected Area. Scientific Reports, 9(1), 10772. https://doi.org/10.1038/s41598-019-47161-0

28. Hutchings, J. A., Ardren, W. R., Barlaup, B. T., Bergman, E., Clarke, K. D., Greenberg, L. A., Lake, C., Piironen, J., Sirois, P., Sundt-Hansen, L. E., & Fraser, D. J. (2019). Life-history variability and conservation status of landlocked Atlantic salmon: An overview. Canadian Journal of Fisheries and Aquatic Sciences, 76(10), 1697–1708. https://doi.org/10.1139/cjfas-2018-0413

29. ICES. (2021). Working Group on North Atlantic Salmon. https://doi.org/10.17895/ICES.PUB.7923

30. Ingólfsson, Ó. A., Einarsson, H. A., & Løkkeborg, S. (2017). The effects of hook and bait sizes on size selectivity and capture efficiency in Icelandic longline fisheries. Fisheries Research, 191, 10–16. https://doi.org/10.1016/j.fishres.2017.02.017

31. Jacobsen, J. & Hansen, L. P. (2001). Feeding habits of wild and escaped farmed Atlantic salmon, *Salmo salar* L., in the Northeast Atlantic. ICES Journal of Marine Science, 58(4), 916–933. https://doi.org/10.1006/jmsc.2001.1084

32. Jensen, A. (1999). Cessation of the Norwegian drift net fishery: Changes observed in Norwegian and Russian populations of Atlantic salmon. ICES Journal of Marine Science, 56(1), 84–95. https://doi.org/10.1006/jmsc.1998.0419

33. Kalinowski, S. T., Manlove, K. R., & Taper, M. L. (2008). ONCOR: A Computer Program for Genetic Stock Identification (Version 2).

34. Kane, A., Ayllón, D., O’Sullivan, R. J., McGinnity, P., & Reed, T. E. (2022). Escalating the conflict? Intersex genetic correlations influence adaptation to environmental change in facultatively migratory populations. Evolutionary Applications, eva.13368. https://doi.org/10.1111/eva.13368

35. Kang, J. koo, & Thibert-Plante, X. (2017). Eco-evolution in size-structured ecosystems: Simulation case study of rapid morphological changes in alewife. BMC Evolutionary Biology, 17(1), 58. https://doi.org/10.1186/s12862-017-0912-4

36. Keefer, M. L., & Caudill, C. C. (2014). Homing and straying by anadromous salmonids: A review of mechanisms and rates. Reviews in Fish Biology and Fisheries, 24(1), 333–368. https://doi.org/10.1007/s11160-013-9334-6

37. Millar, R. B. (1987). Maximum likelihood estimation of mixed stock fishery composition. Canadian Journal of Fisheries and Aquatic Sciences, 44(3), 583–590. https://doi.org/10.1139/f87-071

38. Ó Maoiléidigh, N., White, J., Hansen, L. P., Jacobsen, J. A., Potter, T., Russell, I., & Reddin, D. (2018). Fifty years of marine tag recoveries from Atlantic salmon. ICES Research Report (No. 343; p. 121).

39. Østbye, K., Hagen Hassve, M., Peris Tamayo, A., Hagenlund, M., Vogler, T., & Præbel, K. (2020). “And if you gaze long into an abyss, the abyss gazes also into thee”: Four morphs of Arctic charr adapting to a depth gradient in Lake Tinnsjøen. Evolutionary Applications, 13(6), 1240–1261. https://doi.org/10.1111/eva.12983

40. Otero, J., L’Abée-Lund, J. H., Castro-Santos, T., Leonardsson, K., Storvik, G. O., Jonsson, B., Dempson, B., Russell, I. C., Jensen, A. J., Baglinière, J.-L., Dionne, M., Armstrong, J. D., Romakkaniemi, A., Letcher, B. H., Kocik, J. F., Erkinaro, J., Poole, R., Rogan, G., Lundqvist, H.,… Vøllestad, L. A. (2014). Basin-scale phenology and effects of climate variability on global timing of initial seaward migration of Atlantic salmon (*Salmo salar*). Global Change Biology, 20(1), 61–75. https://doi.org/10.1111/gcb.12363

41. Ozerov, M., Vähä, J.-P., Wennevik, V., Niemelä, E., Svenning, M.-A., Prusov, S., Diaz Fernandez, R., Unneland, L., Vasemägi, A., Falkegård, M., Kalske, T., & Christiansen, B. (2017). Comprehensive microsatellite baseline for genetic stock identification of Atlantic salmon (*Salmo salar L*.) in northernmost Europe. ICES Journal of Marine Science, 74(8), 2159–2169. https://doi.org/10.1093/icesjms/fsx041

42. Ozerov, M., Vasemägi, A., Wennevik, V., Diaz-Fernandez, R., Kent, M., Gilbey, J., Prusov, S., Niemelä, E., & Vähä, J.-P. (2013). Finding Markers That Make a Difference: DNA pooling and SNP-arrays identify population informative markers for genetic stock identification. PLoS ONE, 8(12), e82434. https://doi.org/10.1371/journal.pone.0082434

43. Parrish, D. L., Behnke, R. J., Gephard, S. R., McCormick, S. D., & Reeves, G. H. (1998). Why aren’t there more Atlantic salmon (*Salmo salar*) Canadian Journal of Fisheries and Aquatic Sciences, 55(S1), 281–287. https://doi.org/10.1139/d98-012

44. Pella, J., & Masuda, M. (2008). Bayesian methods for analysis of stock mixtures from genetic characters. Fishery Bulletin, 99(1), 151–167.

45. Potter, E. C. E., Crozier, W. W., Schön, P.-J., Nicholson, M. D., Maxwell, D. L., Prévost, E., Erkinaro, J., Gudbergsson, G., Karlsson, L., Hansen, L. P., MacLean, J. C., Ó Maoiléidigh, N., & Prusov, S. (2004). Estimating and forecasting pre-fishery abundance of Atlantic salmon (*Salmo salar* L.) in the Northeast Atlantic for the management of mixed-stock fisheries. ICES Journal of Marine Science, 61(8), 1359–1369. https://doi.org/10.1016/j.icesjms.2004.08.012

46. R Core Team. (2021). R: A language and environment for statistical computing. https://www.R-project.org/

47. Rikardsen, A. H., & Dempson, J. B. (2010). Dietary Life-Support: The Food and Feeding of Atlantic Salmon at Sea. In Ø. Aas, S. Einum, A. Klemetsen, & J. Skurdal (Eds.), Atlantic Salmon Ecology (pp. 115–143). Wiley-Blackwell. https://doi.org/10.1002/9781444327755.ch5

48. Rikardsen, A. H., Hansen, L. P., Jensen, A. J., Vollen, T., & Finstad, B. (2008). Do Norwegian Atlantic salmon feed in the northern Barents Sea? Tag recoveries from 70 to 78° N. Journal of Fish Biology, 72(7), 1792–1798. https://doi.org/10.1111/j.1095-8649.2008.01823.x

49. Rikardsen, A. H., Righton, D., Strøm, J. F., Thorstad, E. B., Gargan, P., Sheehan, T., Økland, F., Chittenden, C. M., Hedger, R. D., Næsje, T. F., Renkawitz, M., Sturlaugsson, J., Caballero, P., Baktoft, H., Davidsen, J. G., Halttunen, E., Wright, S., Finstad, B., & Aarestrup, K. (2021). Redefining the oceanic distribution of Atlantic salmon. Scientific Reports, 11(1), 12266. https://doi.org/10.1038/s41598-021-91137-y

50. Schindler, D. E., Armstrong, J. B., & Reed, T. E. (2015). The portfolio concept in ecology and evolution. Frontiers in Ecology and the Environment, 13(5), 257–263. https://doi.org/10.1890/140275

51. Spares, A. D., Reader, J. M., Stokesbury, M. J. W., McDermott, T., Zikovsky, L., Avery, T. S., & Dadswell, M. J. (2007). Inferring marine distribution of Canadian and Irish Atlantic salmon (*Salmo salar* L.) in the North Atlantic from tissue concentrations of bio-accumulated caesium 137. ICES Journal of Marine Science, 64(2), 394–404. https://doi.org/10.1093/icesjms/fsl040

52. Strøm, J. F., Rikardsen, A. H., Campana, S. E., Righton, D., Carr, J., Aarestrup, K., Stokesbury, M. J. W., Gargan, P., Javierre, P. C., & Thorstad, E. B. (2019). Ocean predation and mortality of adult Atlantic salmon. Scientific Reports, 9(1), 7890. https://doi.org/10.1038/s41598-019-44041-5

53. Strøm, J. F., Thorstad, E. B., Chafe, G., Sørbye, S. H., Righton, D., Rikardsen, A. H., & Carr, J. (2017). Ocean migration of pop-up satellite archival tagged Atlantic salmon from the Miramichi River in Canada. ICES Journal of Marine Science, 74(5), 1356–1370. https://doi.org/10.1093/icesjms/fsw220

54. Strøm, J. F., Thorstad, E. B., Hedger, R. D., & Rikardsen, A. H. (2018). Revealing the full ocean migration of individual Atlantic salmon. Animal Biotelemetry, 6(1), 2. https://doi.org/10.1186/s40317-018-0146-2

55. Svenning, M.-A., Falkegård, M., Niemelä, E., Vähä, J.-P., Wennevik, V., Ozerov, M., Prusov, S., Dempson, J. B., Power, M., & Fauchald, P. (2019). Coastal migration patterns of the four largest Barents Sea Atlantic salmon stocks inferred using genetic stock identification methods. ICES Journal of Marine Science, 76(6), 1379–1389. https://doi.org/10.1093/icesjms/fsz114

56. Swain, A. (1980). Tagging of salmon smolts in European rivers with special reference to recaptures off West Greenland in 1972 and earlier years (No. 176; Rapports et Procès-Verbaux des Réunions, Conseil Permanent International pour l’Exploration de la Mer, pp. 93–113).

57. Teffer, A. K., Carr, J., Tabata, A., Schulze, A., Bradbury, I., Deschamps, D., Gillis, C.-A., Brunsdon, E. B., Mordecai, G., & Miller, K. M. (2020). A molecular assessment of infectious agents carried by Atlantic salmon at sea and in three eastern Canadian rivers, including aquaculture escapees and North American and European origin wild stocks. FACETS, 5(1), 234–263. https://doi.org/10.1139/facets-2019-0048

58. Thorstad, E. B., Bliss, D., Breau, C., Damon-Randall, K., Sundt-Hansen, L. E., Hatfield, E. M. C., Horsburgh, G., Hansen, H., Maoiléidigh, N. Ó., Sheehan, T., & Sutton, S. G. (2021). Atlantic salmon in a rapidly changing environment—Facing the challenges of reduced marine survival and climate change. Aquatic Conservation: Marine and Freshwater Ecosystems, 31(9), 2654–2665. https://doi.org/10.1002/aqc.3624

59. Thorstad, E. B., Whoriskey, F., Rikardsen, A. H., & Aarestrup, K. (2010). Aquatic Nomads: The Life and Migrations of the Atlantic Salmon. In Ø. Aas, S. Einum, A. Klemetsen, & J. Skurdal (Eds.), Atlantic Salmon Ecology (pp. 1–32). Wiley-Blackwell. https://doi.org/10.1002/9781444327755.ch1

60. Utne, K. R., Pauli, B. D., Haugland, M., Jacobsen, J. A., Maoileidigh, N., Melle, W., Broms, C. T., Nøttestad, L., Holm, M., Thomas, K., & Wennevik, V. (2021). Poor feeding opportunities and reduced condition factor for salmon post-smolts in the Northeast Atlantic Ocean. ICES Journal of Marine Science, 78(8), 2844–2857. https://doi.org/10.1093/icesjms/fsab163

61. Vähä, J.-P., Erkinaro, J., Falkegård, M., Orell, P., & Niemelä, E. (2017). Genetic stock identification of Atlantic salmon and its evaluation in a large population complex. Canadian Journal of Fisheries and Aquatic Sciences, 74(3), 327–338. https://doi.org/10.1139/cjfas-2015-0606

62. Vähä, J.-P., Erkinaro, J., Niemelä, E., Primmer, C. R., Saloniemi, I., Johansen, M., Svenning, M., & Brørs, S. (2011). Temporally stable population-specific differences in run timing of one-sea-winter Atlantic salmon returning to a large river system: Run timing of Atlantic salmon within a river system. Evolutionary Applications, 4(1), 39–53. https://doi.org/10.1111/j.1752-4571.2010.00131.x

63. Vähä, J.-P., Wennevik, V., Ozerov, M., Diaz Fernandez, R., Unneland, L., Haapanen, K., Niemelä, E., Svenning, M.-A., Falkegård, M., Prusov, S., Lyzhov, L., Rysakova, K., Kalske, T., Christiansen, B., & Ustyuzhinsky, G. (2014). Genetic structure of Atlantic salmon in the Barents region and genetic stock identification of coastal fishery catches from Norway and Russia [www.fylkesmannen.no/kolarcticsalmon]. (Kolarctic ENPI CBC — Kolarctic Salmon Project (K0197) Report.).

64. Vihtakari, M. (2022). ggOceanMaps: Plot Data on Oceanographic Maps using ‘ggplot2 (R package version 1.2.6). https://CRAN.R-project.org/package=ggOceanMaps

65. Vollset, K. W., Urdal, K., Utne, K., Thorstad, E. B., Sægrov, H., Raunsgard, A., Skagseth, Ø., Lennox, R. J., Østborg, G. M., Ugedal, O., Jensen, A. J., Bolstad, G. H., & Fiske, P. (2022). Ecological regime shift in the Northeast Atlantic Ocean revealed from the unprecedented reduction in marine growth of Atlantic salmon. Science Advances, 8(9), eabk2542. https://doi.org/10.1126/sciadv.abk2542

