## Supplementary tables and figures for "Genetic stock identification reveals greater use of an oceanic feeding ground around the Faroe Islands by multi-sea winter Atlantic salmon"

Table S1: The population, agglomerative reporting group, and number of Atlantic salmon successfully assigned to a given reporting group used in the analysis. Map number corresponds to positions on Figure 1. The dataset from which each population in the baseline is derived is denoted by superscripts. ^a^ = Bourret et al. 2013; ^b^ = Ozerov et al. 2013; ^c^ = Rikardsen et al. 2004; ^d^ = Erkinaaro & Aykanat unpublished.

| Map number | Population | Reporting group | Individuals assigned and used in analysis | Map number | Population | Reporting group | Individuals assigned and used in analysis |
| --- | --- | --- | --- | --- | --- | --- | --- |
| 1 | Gouffre^a^ | North America | 18 | 26 | Repparfjordelva^b^ | Northern Norway | 0 |
| 2 | Narraguagus^a^ | North America | 0 | 27 | Lakselva^b^ | Northern Norway | 4 |
| 3 | Matapédia^a^ | North America | 41 | 28 | Vestre Jakobselva^b^ | Northern Norway | 1 |
| 4 | Chaloupe^a^ | North America | 17 | 29 | Teno mainstem^d^ | Teno | 24 |
| 5 | Gros Mécatina^a^ | North America | 0 | 30 | Neiden^b^ | Eastern Finnmark, North Kola | 3 |
| 6 | Saint Paul^a^ | North America | 10 | 31 | Titovka^b^ | Eastern Finnmark, North Kola | 0 |
| 7 | Ölfusá^a^ | Iceland | 0 | 32 | Ura^b^ | Eastern Finnmark, North Kola | 2 |
| 8 | Selá^a^ | Iceland | 0 | 33 | Tuloma^a^ | Eastern Finnmark, North Kola | 5 |
| 9 | Moy^a^ | Ireland and UK | 18 | 34 | Kola^b^ | Eastern Finnmark, North Kola | 0 |
| 10 | Blackwater^a^ | Ireland and UK | 66 | 35 | Vindelaiven^a^ | Baltic Sea | 0 |
| 11 | Dionard^a^ | Ireland and UK | 135 | 36 | Torniojoki^a^ | Baltic Sea | 0 |
| 12 | Dart^a^ | Ireland and UK | 55 | 37 | Kunda^a^ | Baltic Sea | 0 |
| 13 | North Esk^a^ | Ireland and UK | 144 | 38 | Narva^b^ | Baltic Sea | 0 |
| 14 | Tweed^a^ | Ireland and UK | 36 | 39 | Pongoma^a^ | Eastern Barents, White Sea | 5 |
| 15 | Narcea^a^ | Bay of Biscay | 0 | 40 | Suma^a^ | Eastern Barents, White Sea | 0 |
| 16 | Piguena^a^ | Bay of Biscay | 0 | 41 | Varzuga Yapoma^a^ | Eastern Barents, White Sea | 1 |
| 17 | Cares^a^ | Bay of Biscay | 0 | 42 | Varzuga mainstem^b^ | Eastern Barents, White Sea | 0 |
| 18 | Loire^a^ | Bay of Biscay | 0 | 43 | Onega^b^ | Eastern Barents, White Sea | 8 |
| 19 | Lardaiselva^a^ | Southern Norway | 113 | 44 | Severnaya Dvina Emtsa^a^ | Eastern Barents, White Sea | 37 |
| 20 | Gaula^a^ | Southern Norway | 592 | 45 | Ponoi mainstem^b^ | Eastern Barents, White Sea | 1 |
| 21 | Numedalslågen^a^ | Southern Norway | 112 | 46 | Ponoi Lebyazhya^a^ | Eastern Barents, White Sea | 9 |
| 22 | Laukhelle^b^ | Northern Norway | 4 | 47 | Mezen Pizhma^b^ | Eastern Barents, White Sea | 1 |
| 23 | Målselv^b^ | Northern Norway | 9 | 48 | Pechora Pizhma^b^ | Eastern Barents, White Sea | 4 |
| 24 | Reisa^b^ | Northern Norway | 2 | 49 | Pechora Unya^b^ | Eastern Barents, White Sea | 44 |
| 25 | Alta^c^ | Northern Norway | 95 | - | - | - | - |

Table S2: Proportion of individual Atlantic salmon correctly assigned to their population of origin and the number of individuals from each baseline population used in the leave-one-out cross-validation of the assignment. Proportion error represented by 95% credible intervals. Percentage of correct and incorrect assignments from leave-one-out cross-validation, as well as population-specific percentages of misidentification are also given. Map number corresponds to positions on Figure 1.

| Map number | Population | Average proportion correctly assigned | 95% credible interval | Number of individuals in baseline used in leave-one-out testing | % Correct | Largest misidentification (Population and percentage) | |
| --- | --- | --- | --- | --- | --- | --- | --- |
| 1 | Gouffre | 0.9986 | (0.9912, 1.0000) | 23 | 95.70 % | Narraguagus | 4.30 % |
| 2 | Narraguagus | 1 | (0.9991, 1.0000) | 19 | 100.00 % | - | - |
| 3 | Matapédia | 0.9897 | (0.9649, 1.0000) | 15 | 93.30 % | Chaloupe | 6.70 % |
| 4 | Chaloupe | 0.9984 | (0.9904, 1.0000) | 12 | 100.00 % | - | - |
| 5 | Gros Mécatina | 0.9997 | (0.9953, 1.0000) | 22 | 100.00 % | - | - |
| 6 | Saint Paul | 0.9989 | (0.9919, 1.0000) | 24 | 95.80 % | Gros Mécatina | 4.20 % |
| 7 | Ölfusá | 1 | (1.0000, 1.0000) | 32 | 96.90 % | Blackwater | 3.10 % |
| 8 | Selá | 1 | (1.0000, 1.0000) | 29 | 100.00 % | - |  |
| 9 | Moy | 0.9799 | (0.9493, 0.9999) | 33 | 81.80 % | Dionard | 12.10 % |
| 10 | Blackwater | 0.9188 | (0.8568, 0.9678) | 37 | 75.70 % | Dart | 8.10 % |
| 11 | Dionard | 0.8649 | (0.8057, 0.9264) | 29 | 51.70 % | North Esk | 20.70 % |
| 12 | Dart | 0.9113 | (0.8488, 0.9641) | 32 | 75.00 % | Blackwater | 21.90 % |
| 13 | North Esk | 0.8798 | (0.8138, 0.9304) | 32 | 68.80 % | Dionard | 18.80 % |
| 14 | Tweed | 0.9341 | (0.8770, 0.9747) | 32 | 62.50 % | Blackwater | 15.60 % |
| 15 | Narcea | 0.9999 | (0.9995, 1.0000) | 15 | 73.30 % | Piguena | 26.70 % |
| 16 | Piguena | 0.9998 | (0.9968, 1.0000) | 12 | 100.00 % | - | - |
| 17 | Cares | 1 | (1.0000, 1.0000) | 27 | 100.00 % | - | - |
| 18 | Loire | 1 | (1.0000, 1.0000) | 34 | 94.10 % | Dart | 5.90 % |
| 19 | Lardaiselva | 0.8828 | (0.8301, 0.9311) | 13 | 76.90 % | Numedalslågen | 7.70 % |
| 20 | Gaula | 0.9809 | (0.9510, 0.9998) | 17 | 94.10 % | Vestre Jakobselv | 5.90 % |
| 21 | Numedalslågen | 0.9967 | (0.9840, 1.0000) | 4 | 100.00 % | - | - |
| 22 | Laukhelle | 0.9931 | (0.9803, 1.0000) | 40 | 92.50 % | Gaula | 2.50 % |
| 23 | Målselv | 0.9922 | (0.9816, 1.0000) | 40 | 85.00 % | Gaula | 2.50 % |
| 24 | Reisa | 0.996 | (0.9836, 1.0000) | 40 | 97.50 % | Repparfjordelva | 2.50 % |
| 25 | Alta | 0.9988 | (0.9925, 1.0000) | 62 | 90.30 % | Gaula | 3.20 % |
| 26 | Repparfjordelva | 0.953 | (0.9173, 0.9873) | 40 | 70.00 % | Titovka | 10.00 % |
| 27 | Lakselva | 0.9958 | (0.9814, 1.0000) | 40 | 95.00 % | Gaula | 2.50 % |
| 28 | Vestre Jakobselva | 0.9682 | (0.9390, 0.9936) | 40 | 77.50 % | Repparfjordelva | 7.50 % |
| 29 | Teno mainstem | 0.998 | (0.9904, 1.0000) | 279 | 91.80 % | Alta | 2.90 % |
| 30 | Neiden | 0.9701 | (0.9400, 0.9948) | 40 | 77.50 % | Titovka | 10.00 % |
| 31 | Titovka | 0.8672 | (0.7984, 0.9158) | 40 | 60.00 % | Ura | 22.50 % |
| 32 | Ura | 0.9006 | (0.8378, 0.9480) | 40 | 72.50 % | Titovka | 22.50 % |
| 33 | Tuloma | 0.9865 | (0.9642, 0.9999) | 35 | 85.70 % | Titovka | 8.60 % |
| 34 | Kola | 1 | (1.0000, 1.0000) | 87 | 79.30 % | Yapoma | 4.60 % |
| 35 | Vindelaiven | 1 | (1.0000, 1.0000) | 33 | 100.00 % | - | - |
| 36 | Torniojoki | 0.9999 | (0.9962, 1.0000) | 27 | 100.00 % | - | - |
| 37 | Kunda | 1 | (1.0000, 1.0000) | 37 | 94.60 % | Narva | 5.40 % |
| 38 | Narva | 1 | (1.0000, 1.0000) | 40 | 100.00 % | - | - |
| 39 | Pongoma | 1 | (1.0000, 1.0000) | 32 | 96.90 % | Yapoma | 3.10 % |
| 40 | Suma | 1 | (1.0000, 1.0000) | 23 | 100.00 % | - | - |
| 41 | Varzuga Yapoma | 0.8875 | (0.8209, 0.9466) | 35 | 77.10 % | Varzuga mainstem | 14.30 % |
| 42 | Varzuga mainstem | 0.9225 | (0.8749, 0.9620) | 40 | 80.00 % | Varzuga Yapoma | 17.50 % |
| 43 | Onega | 1 | (1.0000, 1.0000) | 40 | 100.00 % | - | - |
| 44 | Severnaya Dvina Emtsa | 0.9997 | (0.9954, 1.0000) | 37 | 91.90 % | Teno | 5.40 % |
| 45 | Ponoi mainstem | 0.9654 | (0.9276, 0.9916) | 40 | 77.50 % | Lebyazhya | 10.00 % |
| 46 | Ponoi Lebyazhya | 0.9562 | (0.9180, 0.9883) | 35 | 82.90 % | Ponoi mainstem | 14.30 % |
| 47 | Mezen Pizhma | 1 | (0.9997, 1.0000) | 40 | 100.00 % | - | - |
| 48 | Pechora Pizhma | 0.9999 | (1.0000, 1.0000) | 40 | 100.00 % | - | - |
| 49 | Pechora Unya | 1 | (1.0000, 1.0000) | 40 | 100.00 % | - | - |

Table S3: Proportion of individual Atlantic salmon correctly assigned to their agglomerative reporting group and the number of individuals from each reporting group used in the leave-one-out cross-validation of the assignment. Percentage of correct and incorrect assignments from leave-one-out cross-validation, as well as group-specific percentages of misidentification are also given.

| Reporting group | Average proportion correctly assigned | 95% credible interval | Number of individuals in baseline used in leave-one-out testing | % Correct | Largest misidentification (Reporting group and percentage) |
| --- | --- | --- | --- | --- | --- |
| North America | 1 | (1.0000, 1.0000) | 23 | 100.00 % |  |
| North America | 1 | (1.0000, 1.0000) | 19 | 100.00 % | - |
| North America | 1 | (1.0000, 1.0000) | 15 | 100.00 % | - |
| North America | 1 | (1.0000, 1.0000) | 12 | 100.00 % | - |
| North America | 1 | (1.0000, 1.0000) | 22 | 100.00 % | - |
| North America | 1 | (1.0000, 1.0000) | 24 | 100.00 % | - |
| Iceland | 1 | (1.0000, 1.0000) | 32 | 96.90 % | - |
| Iceland | 1 | (1.0000, 1.0000) | 29 | 100.00 % | - |
| Ireland and UK | 0.9998 | (0.9986, 1.0000) | 33 | 97.00 % | - |
| Ireland and UK | 0.9997 | (0.9954, 1.0000) | 37 | 100.00 % | - |
| Ireland and UK | 0.9993 | (0.9935, 1.0000) | 29 | 100.00 % | - |
| Ireland and UK | 0.9999 | (1.0000, 1.0000) | 32 | 100.00 % | - |
| Ireland and UK | 0.9996 | (0.9951, 1.0000) | 32 | 100.00 % | - |
| Ireland and UK | 1 | (1.0000, 1.0000) | 32 | 100.00 % | - |
| Bay of Biscay | 1 | (1.0000, 1.0000) | 15 | 100.00 % | - |
| Bay of Biscay | 1 | (1.0000, 1.0000) | 12 | 100.00 % | - |
| Bay of Biscay | 1 | (1.0000, 1.0000) | 27 | 100.00 % | - |
| Bay of Biscay | 1 | (1.0000, 1.0000) | 34 | 94.10 % | - |
| Southern Norway | 0.9929 | (0.9760, 1.0000) | 13 | 92.30 % | - |
| Southern Norway | 0.9953 | (0.9817, 1.0000) | 17 | 94.10 % | - |
| Southern Norway | 0.9991 | (0.9944, 1.0000) | 4 | 100.00 % | - |
| Northern Norway | 0.9963 | (0.9855, 1.0000) | 40 | 97.50 % | - |
| Northern Norway | 0.9971 | (0.9863, 1.0000) | 40 | 97.50 % | - |
| Northern Norway | 0.9999 | (0.9999, 1.0000) | 40 | 100.00 % | - |
| Northern Norway | 0.9996 | (0.9943, 1.0000) | 62 | 91.90 % | - |
| Northern Norway | 0.9772 | (0.9510, 0.9975) | 40 | 82.50 % | - |
| Northern Norway | 0.9992 | (0.9931, 1.0000) | 40 | 97.50 % | - |
| Northern Norway | 0.9885 | (0.9643, 1.0000) | 40 | 92.50 % | - |
| Teno | 0.998 | (0.9904, 1.0000) | 279 | 91.80 % | Eastern Finnmark, North Kola 2.90% |
| Eastern Finnmark, North Kola | 0.9939 | (0.9798, 1.0000) | 40 | 92.50 % | - |
| Eastern Finnmark, North Kola | 0.9834 | (0.9569, 1.0000) | 40 | 87.50 % | - |
| Eastern Finnmark, North Kola | 0.9975 | (0.9862, 1.0000) | 40 | 95.00 % | - |
| Eastern Finnmark, North Kola | 0.9927 | (0.9778, 1.0000) | 35 | 100.00 % | - |
| Eastern Finnmark, North Kola | 1 | (1.0000, 1.0000) | 87 | 79.30 % | - |
| Baltic Sea | 1 | (1.0000, 1.0000) | 33 | 100.00 % | - |
| Baltic Sea | 1 | (1.0000, 1.0000) | 27 | 100.00 % | - |
| Baltic Sea | 1 | (1.0000, 1.0000) | 37 | 100.00 % | - |
| Baltic Sea | 1 | (1.0000, 1.0000) | 40 | 100.00 % | - |
| Eastern Barents, White Sea | 1 | (1.0000, 1.0000) | 32 | 100.00 % | - |
| Eastern Barents, White Sea | 1 | (1.0000, 1.0000) | 23 | 100.00 % | - |
| Eastern Barents, White Sea | 0.9999 | (0.9970, 1.0000) | 35 | 97.10 % | - |
| Eastern Barents, White Sea | 1 | (1.0000, 1.0000) | 40 | 100.00 % | - |
| Eastern Barents, White Sea | 1 | (1.0000, 1.0000) | 40 | 100.00 % | - |
| Eastern Barents, White Sea | 1 | (1.0000, 1.0000) | 37 | 91.90 % | Eastern Finnmark, North Kola 2.70% |
| Eastern Barents, White Sea | 0.9999 | (0.9978, 1.0000) | 40 | 100.00 % | - |
| Eastern Barents, White Sea | 0.9996 | (0.9955, 1.0000) | 35 | 100.00 % | - |
| Eastern Barents, White Sea | 1 | (1.0000, 1.0000) | 40 | 100.00 % | - |
| Eastern Barents, White Sea | 1 | (1.0000, 1.0000) | 40 | 100.00 % | - |
| Eastern Barents, White Sea | 1 | (1.0000, 1.0000) | 40 | 100.00 % | - |

Table S4: Reported catch in number of fish of small (<3 kg) medium (3-7 kg) and large salmon (>7 kg). The small, medium and large salmon are assumed to be 1SW, 2SW, and 3SW salmon, respectively.

| ICES region | Year | Small | Medium | Large |
| --- | --- | --- | --- | --- |
| Norway south-east | 1993 | 23070 | 10689 | 1907 |
| Norway south-east | 1994 | 23987 | 8468 | 1520 |
| Norway south-east | 1995 | 21847 | 9864 | 1766 |
| Norway south-east | 1996 | 20738 | 11366 | 2172 |
| Norway south-west | 1993 | 11433 | 7338 | 2901 |
| Norway south-west | 1994 | 18597 | 8596 | 2365 |
| Norway south-west | 1995 | 10863 | 10954 | 2168 |
| Norway south-west | 1996 | 7048 | 9727 | 2819 |
| Norway middle | 1993 | 58338 | 18079 | 10105 |
| Norway middle | 1994 | 113426 | 24792 | 8728 |
| Norway middle | 1995 | 57823 | 32777 | 9920 |
| Norway middle | 1996 | 28936 | 20111 | 11502 |
| Norway north | 1993 | 42610 | 20178 | 12769 |
| Norway north | 1994 | 36117 | 20413 | 9137 |
| Norway north | 1995 | 34223 | 15391 | 8959 |
| Norway north | 1996 | 42794 | 24399 | 7974 |

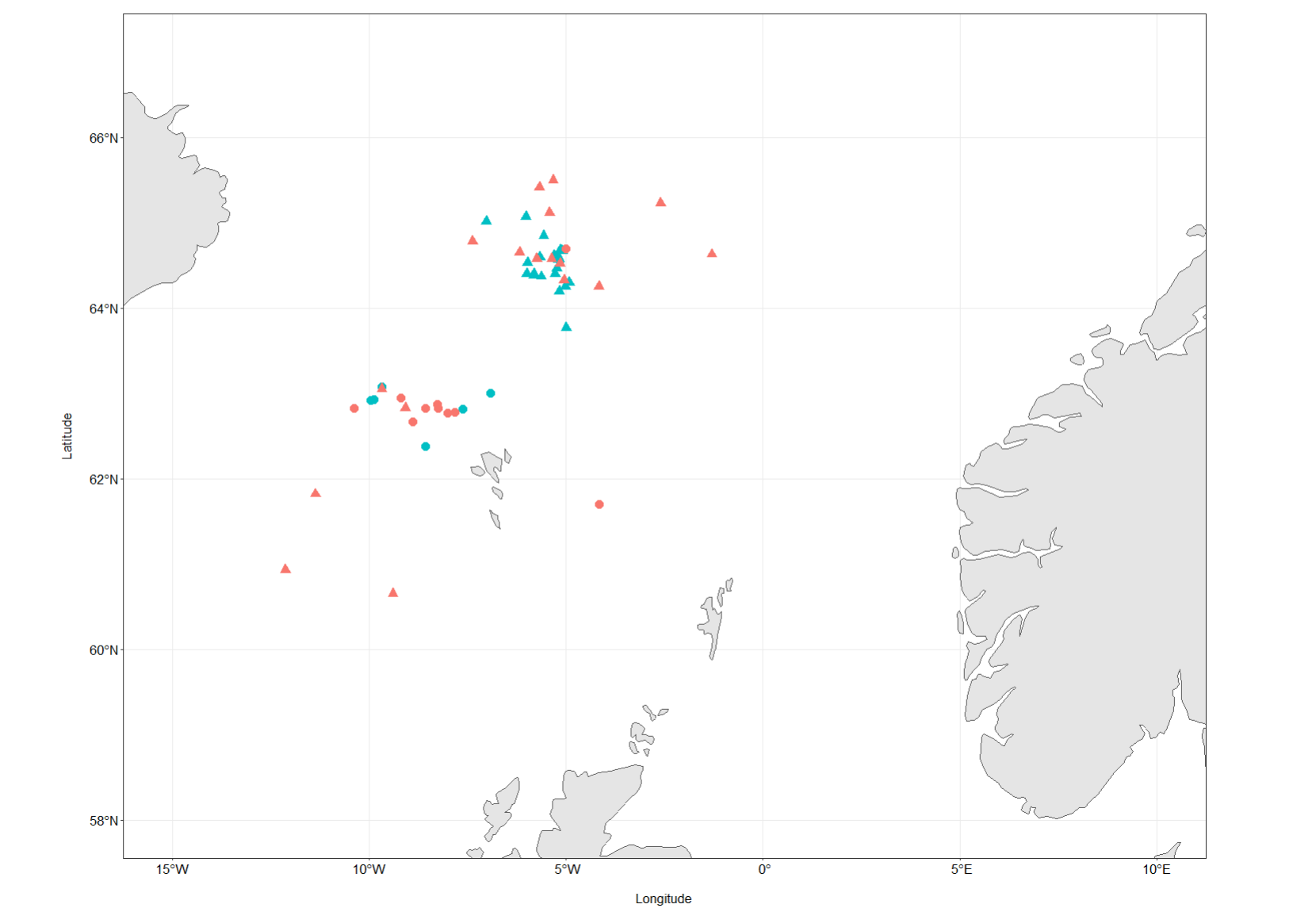

Figure S1: Location, season, and fishing period of each of the experimental longlines used to sample Atlantic salmon from their oceanic feeding ground around the Faroe Islands.

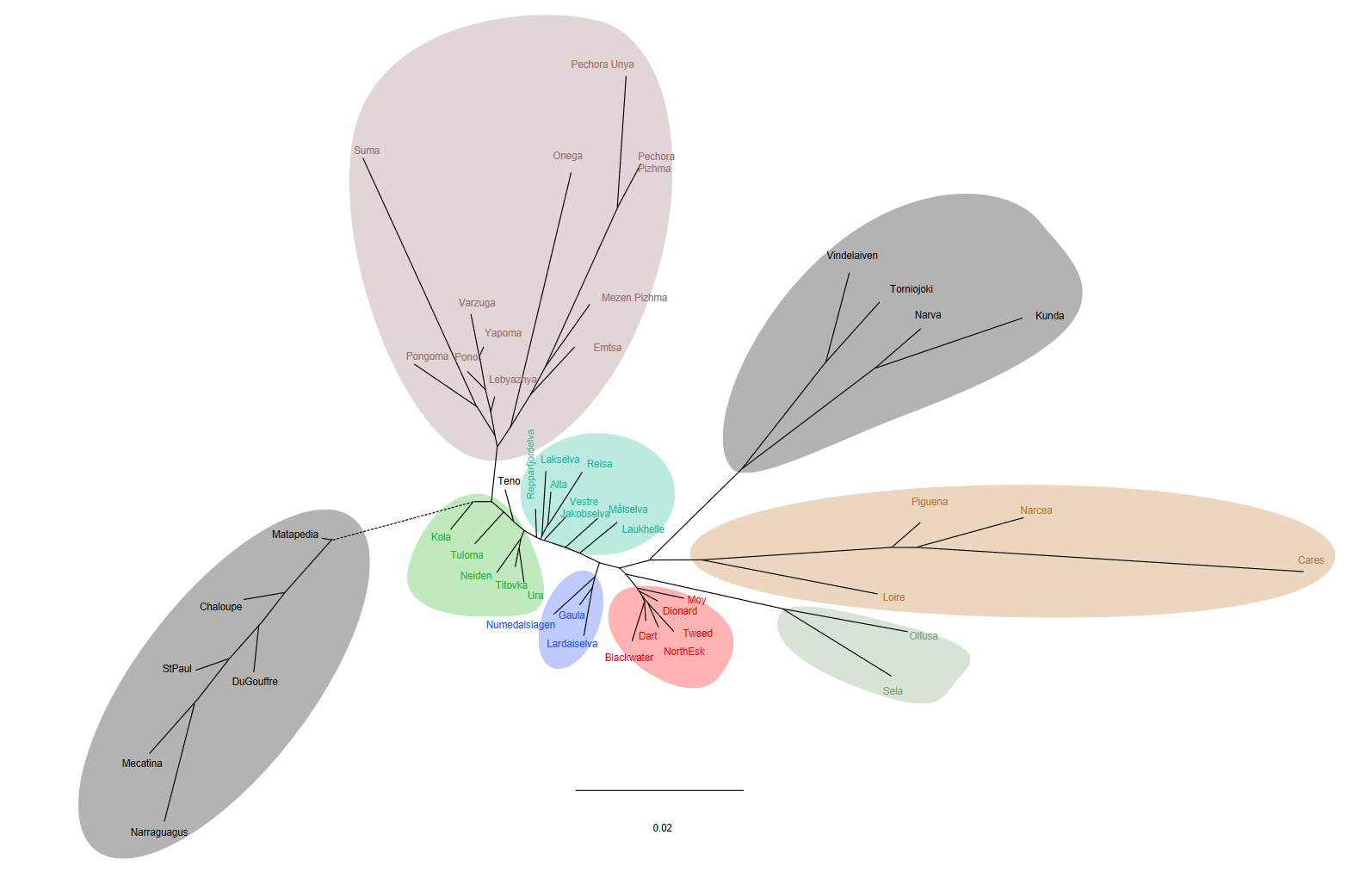

Figure S2: Neighbour joining tree used to inform the delineation of agglomerative reporting groups.
